# Targeting D-Ribose-Binding Proteins in *Brucella melitensis*: A Novel Frontier Against Antibiotic Resistance

**DOI:** 10.1101/2025.02.28.640864

**Authors:** Omid Moradi, Ali Maghsoudi, Ali Akbar Masoudi, Rasoul Vaez Torshizi

## Abstract

**Background:** Antibiotic resistance among pathogens common to human beings and animals, which include *Brucella melitensis*, has end up a significant worldwide health task. Traditional antibiotic treatments for brucellosis, along with lengthy-time period regimens of doxycycline and rifampicin, are going through increasing boundaries because of rising resistance, affected person adherence issues, and considerable side results.

**Methods:** This observe investigates the capacity of targeting the periplasmic D-ribose-binding protein (DBP), a key component of the bacterial ATP-binding cassette (ABC) delivery system, as a unique healing technique. Protein structural modeling was performed to use of superior computational tools together with AlphaFold, Swiss-Model, and Phyre2, followed by validation via Ramachandran plots and energy minimization techniques. Molecular docking analyses recognized D-Talopyranose as a promising ligand with a high binding affinity of -5.8 kcal/mol. Subsequent ADMET profiling found out favorable pharmacokinetic and toxicological results, assisting its potential as a drug candidate. Molecular dynamics simulations similarly evaluated the stability and dynamics of the protein-ligand interplay complex, confirming its suitability for therapeutic programs.

**Results:** Advanced computational tools were used to analyze the protein’s structure, and key modifications that influence its stability and function were identified. AlphaFold was recognized as the most reliable model for predicting the protein’s 3D architecture, with its predictions being validated by metrics such as Ramachandran plots and error assessments. D-Talopyranose, a sugar molecule, was revealed as a top candidate through molecular docking due to its strong binding affinity with DBP. Promising drug-like properties, including balanced solubility, low toxicity risks, and minimal interactions with metabolic enzymes, were highlighted in further analysis, though environmental concerns around biodegradability were noted. The stability of the protein-ligand complex was tracked through simulations, showing consistent structural integrity despite minor flexibility in certain regions. While frequent dosing may be required due to the compound’s rapid clearance from the body, its safety profile and synthetic accessibility are positioned as a viable starting point for drug development. Computational predictions are bridged with practical insights in this work, offering a roadmap for targeted therapies against antibiotic-resistant infections, while the need to balance efficacy with environmental safety in future optimizations is underscored. Our outcomes reveal that targeting DBP could offer a unique mechanism to combat antibiotic-resistant lines of *Brucella melitensis* by using disrupting essential metabolic pathways.

**Conclusion:** This study affords a promising street for revolutionary brucellosis treatments by way of addressing the challenges posed using antibiotic resistance and paves the manner for experimental validation and optimization of the identified ligands. Such focused strategies may also notably improve ailment control and reduce the worldwide burden of brucellosis, mainly in areas where traditional antibiotics are losing their efficacy.

## 1. Introduction

Zoonoses are diseases that can be transmitted from animals to humans. They can be caused by various pathogens, including bacteria, viruses, parasites, and fungi. These diseases can spread through direct contact with animals, consumption of contaminated food/water, or through vectors like mosquitoes and parasites. Some common examples of zoonotic diseases include Rabies, Lyme disease, Avian Influenza and Brucellosis. Brucellosis is a specific type of zoonosis caused by bacteria of the genus Brucella. It primarily affects livestock such as cattle, goat, and sheep, but can also infect humans [1]. The Brucella bacteria cause this disease, which remains one of the most common zoonotic infections [2]. The World Organization for Animal Health (WOAH) views brucellosis as a significant concern due to its impacts on public health, economic losses, and disruptions to international trade [3].

This disease burdens livestock with substantial costs, as different Brucella species infect various animals: cows (*B. abortus*), goats (*B. melitensis*), dogs (*B. canis*), sheep (*B. ovis*), pigs (*B. suis*), and rodents (*B. neotomae*). The broad range of affected species underscores the widespread impact of brucellosis [4]. Brucellosis is prevalent among people working closely with animals, like farmers and veterinarians, due to exposure to infected animal secretions [5]. Individuals can contract the disease by handling infected animals, consuming contaminated livestock products, or inhaling the bacteria [6]. Human symptoms of brucellosis include fever, sweats, malaise, anorexia, headache, and muscle pain. However, chronic brucellosis can result in severe complications such as arthritis, endocarditis, and neurological disorders [7]. The disease starts as an acute infection and can progress to a chronic condition with various complications. Annually, around 500,000 new cases are estimated, but this number may be underestimated due to diagnostic challenges, especially in areas with limited healthcare [8].

The ATP-binding cassette (ABC) transport system is essential for transporting various molecules across cellular membranes in all species, including bacteria, archaea, and eukaryotes, using energy from ATP hydrolysis. The transport process involves substrate binding, ATP hydrolysis at nucleotide binding domains, conformational changes, substrate translocation, release into the cytoplasm, resetting for another cycle. ABC transporters handle substrates like nutrients, ions, drugs, toxins, and lipids. In bacteria, these transporters are crucial for nutrient uptake and can dismiss toxic compounds, contributing to antibiotic resistance. They also play roles in pathogen virulence by secreting virulence factors and in metabolic regulation by maintaining nutrient levels, highlighting their importance in cellular processes across diverse organisms [9-12].

The DBP is essential in the bacterial ABC transport system for D-Ribose uptake, an essential sugar for bacterial metabolism. It binds and transports D-Ribose across the bacterial cell membrane into the cytoplasm. This protein specifically interacts with D-Ribose in the periplasmic space to facilitate its membrane transport [13]. This process is crucial for bacterial energy metabolism, converting D-Ribose to ATP necessary for cellular activities. Binding D-Ribose causes the protein to change shape, aiding interaction with the ABC transport system for efficient transport. Transporting and metabolizing D-Ribose is crucial for bacterial survival and growth, especially in sugar-rich environments. This nutrient absorption illustrates how bacteria adapt and optimize their metabolism for energy generation [9-11, 14, 15].

Brucellosis is usually treated with a combination of antibiotics, such as doxycycline and rifampicin, often alongside an aminoglycoside, over several weeks to months. However, prolonged treatment presents challenges, including difficulties in patient adherence, increased side effects like gastrointestinal issues and liver toxicity, and the emergence of antibiotic-resistant strains, complicating future treatments [11]. The financial burden on patients and healthcare systems also rises due to the long-term medication and managing side effects. Thus, while lengthy treatment is essential for completely eradicating Brucella bacteria, it significantly affects patients’ quality of life and presents several challenges that need addressing. The DBP in Brucella bacteria is critical for drug discovery and understanding bacterial metabolism. *Brucella melitensis*, which is intracellular pathogen, utilizes sugars like D-Ribose for survival and replication within host macrophages, key for energy metabolism and pathogenicity. Targeting this protein with specific drugs could reduce off-target effects and side effects, disrupt bacterial metabolism, and potentially lead to bacterial cell death, providing a new mechanism of action against infections, especially amidst rising antibiotic resistance [5, 16-18].

With traditional antibiotics losing their effectiveness, targeting novel proteins like the DBP presents promising alternatives. By studying its structure and function, new drugs can be developed to inhibit its activity, offering effective treatments for antibiotic-resistant bacterial infections. For instance, research on Epigallocatechin Gallate (EGCG) from green tea shows it targets key bacterial metabolism proteins, providing insights into potential drug mechanisms against resistant strains like *Mycobacterium Tuberculosis* [19]. The discovery of small molecules that bind to viral proteins emphasizes the significance of targeting specific binding sites in drug discovery, similar to targeting DBPs. The creation of inhibitors for Poly Glycohydrolase showcases the value of exploring new mechanisms in drug development, paralleling the approach of targeting DBPs in bacteria. These studies highlight the crucial role of specificity and innovative mechanisms in developing new therapeutics, demonstrating how targeting specific bacterial proteins can result in effective treatments for infections [20-22]. The capability of Brucella to obtain essential nutrients, such as iron, during macrophage infection is crucial for its survival, indicating that nutrient transport systems, including those for sugars like D-Ribose, are crucial to its pathogenic strategy [23-27]. This study aims to concentrating on the DBP to identify new therapeutic strategies to hinder Brucella’s virulence and pathogenicity. Inhibiting the DBP could disrupt essential bacterial metabolic pathways, thus diminishing their infectious capabilities. This method also presents an alternative to traditional antibiotics, tackling the increasing issue of antibiotic resistance and offering a novel approach for effective brucellosis treatments.

## 2. Material And Methods

### 2.1. Data retrieval

The protein sequence of the DBP was retrieved from *Brucella melitensis* biotype 1, as documented in UniProt entry Q8YCU3 (Q8YCU3_BRUME), corresponding to strain ATCC 23456 / CCUG 17765 / NCTC 10094 / 16M.

### 2.2. Gene Ontology

The DBP precursor from *Brucella melitensis* biotype 1, identified in UniProt under entry Q8YCU3, is a notable molecular structure. This protein, which consists of 292 amino acids, is situated in the periplasmic space of the bacterium. It plays an essential role in nutrient absorption and cellular metabolism. It’s functional properties include carbohydrate binding and amino acid transport, which are critical for the organism’s survival and operation [28]. This similarity underscores its potential binding affinity and specificity for ribose.

### 2.3. 3D Structure Prediction

Six 3D Structure models of the sequence using various computational tools, including Swiss-Model server [29], AlphaFold Colab server [30], Robetta server [31], I-Tasser server [32], Phyre2 server [33], and C-Quark server [34] was generated. Subsequently, all 3D structures were subjected to energy minimization using the Yasara web server [35]. Following this, polar hydrogens were added, and Kollman charges were assigned using AutoDock Tools software [36]. The Saves server offers several tools for validating and analyzing protein structures, The analysis includes Procheck, which evaluates several key factors. The Ramachandran Plot ensures the torsional angles of amino acid residues fall within permitted regions, indicating stable and functional conformations. The overall G-factors evaluation confirms the structural integrity of the protein model, leading to more accurate and reliable predictions. Additionally, checking planar groups to ensure they are within limits show that most planar groups meet expected geometric standards, thereby enhancing the structural integrity, reliability of predictions, confidence in results, and overall optimization [37, 38].

To identify potential errors within the protein structure, ERRAT analysis was used for the data of non-bonded interactions between various atom types. This analysis is crucial for ensuring the accuracy of the protein model, which is essential in drug discovery [39]. The Verify 3D evaluation produced the following outcomes for the averaged 3D-1D scores of the residues, demonstrating how well the residues align with the predicted three-dimensional structure of the protein. This evaluation is crucial for confirming the protein model’s accuracy and reliability, which is crucial for its application in drug discovery and other fields [40, 41].

QMEAN was calculated by the SWISS-MODEL webserver [42], and the Z-score was calculated by the ProSA-web server [43, 44]. These metrics are essential for evaluating the quality and reliability of protein models. QMEAN delivers a composite score evaluating various structural features, aiding in the identification of potential errors and ensuring the model’s overall quality [42]. ProSA-web server Z-score shows how well the model matches typical structures of a similar size, pointing out any major deviations from expected norms [43, 44]. These tools offer a thorough assessment of the protein model’s accuracy and reliability, which is crucial for its use in drug discovery and structural biology. The Global_score was determined using the VOROMQA server (Voronoi-based Model Quality Assessment), which evaluates protein models by analyzing the spatial arrangement of atoms through Voronoi tessellation. This global score is obtained by summing the local scores calculated for each residue in the protein structure [45].

The optimal 3D structure was identified and selected based on average metrics using a Python script (supplementary 2). This script imported and converted metric columns to numeric values, calculated the average score for each model while ignoring NaN (Not a Number) values, and determined the model with the highest average score as the best. The dataset included structures from AlphaFold, Swiss-Model, I-Tasser, Phyre2, C-Quark, and Robetta.

after that pLDDT and PAE Analyses were used to assess the reliability and accuracy of the predicted protein structure. The pLDDT scores provided a residue-by-residue confidence level, with higher values indicating regions of the model where the predicted 3D positions were most trustworthy. Meanwhile, the PAE heatmap visualized alignment errors between residue pairs, where darker shades reflected lower errors and stronger spatial consistency across the structure. Together, these metrics validated the structural coherence of the model—widespread high pLDDT scores and low PAE values suggested that the overall architecture was well-predicted, with minimal local or global uncertainties. This dual analysis ensure that the model’s confidence aligned with experimental expectations, making it a robust foundation for further functional studies or drug discovery efforts.

### 2.4. PTM (Post Translation Modification) Analysis

Extensive analysis of the protein sequence to identify potential post-translational modifications (PTMs) and evaluate various physicochemical properties was conducted. To facilitate this process, the computational environment was set up, and essential libraries such as Biopython [46], Matplotlib [47], and NumPy [48] were used. primary objective was to identify potential phosphorylation sites within the protein sequence. Phosphorylation plays a crucial role in regulating protein functions by modifying activity and interactions. Using regular expressions, the sequence was systematically scanned to locate serine (S), threonine (T), and tyrosine (Y) residues. These residues were marked as potential sites for phosphorylation, highlighting key regions that may play pivotal roles in the protein’s regulatory mechanisms. Next, the sequence was assessed for N-terminal acetylation, a common modification if it starts with methionine (M). N-glycosylation motifs were also scanned for in the sequence. These motifs indicate sites for glycan attachment, which can impact protein folding and stability. Then, various physicochemical properties of the cleaned and modified protein sequence were calculated. Hydrophobicity was assessed using the Kyte-Doolittle scale [49], the instability index was calculated to predict protein stability, and the molecular weight was determined. Additionally, the isoelectric point was estimated to understand the protein’s solubility under different pH levels. The aliphatic index was calculated to evaluate thermostability, and aromaticity was assessed by counting aromatic residues (F, Y, and W).

These analyses provided a holistic understanding of the protein’s structural and functional characteristics. They offered valuable insights into the protein’s behavior and potential interactions within a biological context, which could be critical for further research and therapeutic development.

### 2.5. Active Site Prediction

A precise prediction for the active site was obtained using Molegro Virtual Docker version 6 (Figure 1). This software enabled the precise identification of potential binding sites [50]. To ensure we captured all meaningful cavities, we set a minimum cavity size of 10 cubic angstroms. Using a probe size of 2 angstroms, we carefully mapped out the binding site landscape to pinpoint the most likely active sites. These carefully chosen parameters not only improve the accuracy of our predictions but also lay a strong groundwork for developing new drugs in the future.

**Figure 1.**
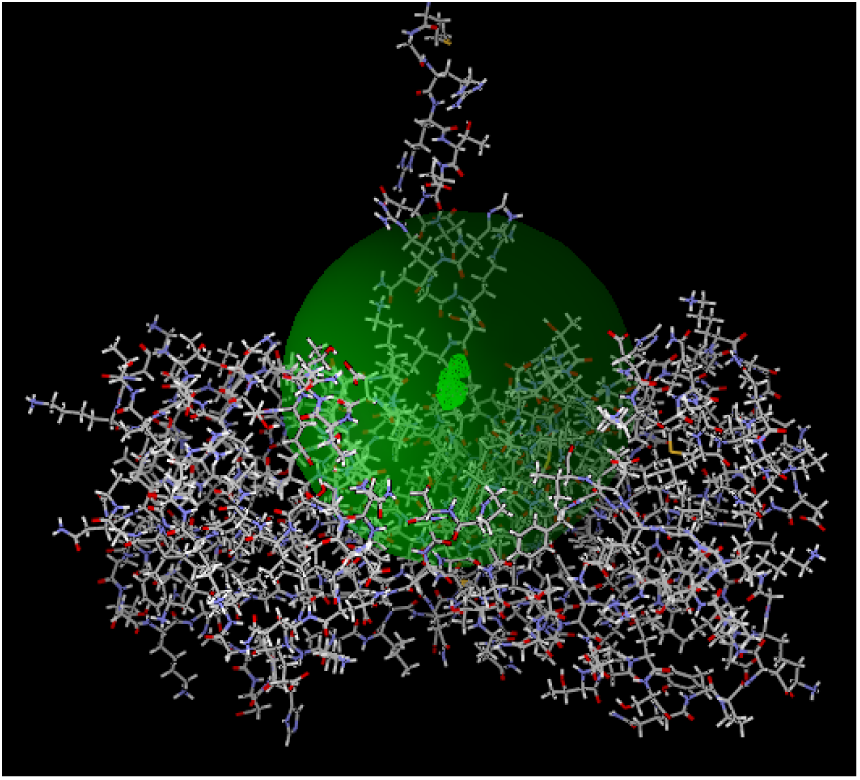
Visualization of the predicted active site in a AlphaFold structure, highlighted by a green sphere. Provided by Molegro 6 software.

### 2.6. Ligand Preparation

D-Ribose (CID 10975657) was used as the specific ligand and control compound, making it the benchmark for comparison. One hundred compounds were retrieved from the PubChem database to identify similar interactions. A Tanimoto similarity threshold of 90% was employed, ensuring that these compounds had at least 90% structural similarity to D-Ribose, thereby guaranteeing comparable binding properties. Once the compounds were obtained, they were converted from their initial SDF format to PDBQT format for further computational analysis. During this conversion process, MMFF energy minimization [51] was applied to each compound. This step refined the molecular structure, ensuring each compound was in its most stable conformation, which was crucial for accurate docking simulations. OpenBabel 2.4.1 software was used for this process [52].

### 2.7. Molecular Docking

Docking simulations of the protein structure were conducted with 100 compounds retrieved from the PubChem database. AutoDock Vina through PYRx - Python Prescription 0.8 software was used for the docking analysis [53]. This high-throughput approach allowed for efficient screening of potential ligands. Following the docking simulations, the results were meticulously analyzed and sorted based on binding affinity. The compounds with the strongest potential interactions with the protein were identified. This process was crucial for pinpointing candidates with the highest likelihood of effective binding, providing a solid foundation for subsequent experimental validation and drug discovery efforts [54].

### 2.8. ADMET (Absorption, Distribution, Metabolism, Excretion, and Toxicity) Analysis

ADMET analysis, which encompasses Absorption, Distribution, Metabolism, Excretion, and Toxicity, is a fundamental aspect of drug development. This evaluation allows us to examine the pharmacokinetic and toxicological profiles of potential drug candidates, ensuring their safety and efficacy before they reach the clinical trial stage [55]. Evaluating the promising compound helps focus the candidate features for interaction with body in future analyses, thereby reducing high attrition rates in drug development. In our analysis, the ADMETlab 2 webserver was utilized to gain comprehensive insights into the pharmacokinetic and toxicological profiles of D-Talose, providing valuable data for making informed decisions about its development [56].

### 2.9. Molecular Dynamics Simulation

GROMACS-2021.4 was used to conduct molecular dynamic simulations of the target protein and its best protein-ligand complexes [57]. The initial structure of the protein was prepared using the CHARMM27 force field and the TIP3P water model[58]. The protein-ligand complex was then enclosed in a triclinic box with a 1.0 nm buffer distance from the edges and solvated using the SPC216 water model. The system was ionized by adding Na^+^ and Cl^−^ ions to reach a concentration of 0.1 M, ensuring overall neutrality [59, 60]. Subsequently, energy minimization was conducted to eliminate bad contacts within the system. Equilibration steps were performed in two phases: NVT (constant Number of particles, Volume, and Temperature) followed by NPT (constant Number of particles, Pressure, and Temperature) using predefined molecular dynamics parameters. The production molecular dynamics run was executed for an extended simulation time to obtain stable conformations and reliable interaction data [61-64]. Once the simulation was completed, we re-centered and re-wrapped the trajectory to ensure accurate visualization and analysis, including RMSD, RMSF, hydrogen bond calculations, radius of gyration, and energy calculations, were then performed to evaluate the stability and interactions within the protein-ligand complexes. The results were visualized using Xmgrace, which allows to examine and confirm the binding affinities and interactions at the molecular level in great detail [65, 66].

## 3. Results And Discussion

### 3.1. Post Translation Modification

The PTM (Post-Translational Modifications) analysis identified several key modifications in the protein sequence. There are potential phosphorylation sites at numerous positions, including 5, 13, 22, 28, 34, 45, 47, 49, 54, 57, 60, 73, 106, 117, 122, 129, 137, 143, 145, 148, 150, 153, 155, 171, 192, 222, 236, 237, 246, 259, 262, 277, 284, and 286. Additionally, there is a potential N-terminal acetylation and an N-glycosylation site at position 58. The analysis also provided hydrophobicity (Kyte-Doolittle scale) at -0.397, an instability index of 54.49, a molecular weight of approximately 34,233.28 Da, an isoelectric point of 5.60, an aliphatic index of 31.90, and an aromaticity value of 0.046. Overall, these modifications and metrics provide a detailed insight into the stability, structure, and potential functional sites within the protein.

### 3.2. 3D structure Preparation and Validation

The prepared 3d structures was evaluated by several ways, including Ramachandran Plot, ERRAT, verify 3d, overall G-Factors, Planar Groups, Qmean, Global-Score, and Z-Score (Table 1). The optimal 3D structure was selected based on average metrics calculated using a Python script. This script imported data, converted metric columns into numeric values, and computed the average score for each model, ignoring NaN values. The model with the highest average score was chosen as the best. The dataset included models from AlphaFold, Swiss-model, I-Tasser, Phyre2, C-Quark, and Robetta servers.

**Table 1.** Comprehensive Evaluation Metrics for 3D Protein Structures Generated by Different Tools.

|  | Ramachandran Favored | ERRAT | Verify 3D | Overall G-factors | Planar Groups | Qmean | Global_Score | z-score |
| --- | --- | --- | --- | --- | --- | --- | --- | --- |
| **AlphaFold** | 93.70% | 98.9091 | 69.86 | 0.22 | 93.90% | 0.26 | 0.456626 | −8.56 |
| **Swiss-Model** | 92.80% | 97.2441 | 67.40 | 0.19 | 96.60% | −0.16 | 0.442839 | −8.11 |
| **I-Tasser** | 90.60% | 96.0854 | 68.49 | 0.09 | 93.90% | −1.94 | 0.415498 | −7.47 |
| **Phyre2** | 92.10% | 99.6154 | 64.86 | 0.22 | 92.00% | −0.52 | 0.444604 | −7.98 |
| **C-Quark** | 88.20% | 97.8495 | 65.75 | 0.04 | 93.90% | −2.33 | 0.401066 | −8.01 |
| **Robetta** | 92.50% | 95.4064 | 71.23 | 0.24 | 93.90% | 0.68 | 0.452642 | −8.20 |

Table 1 presents the evaluation metrics for 3D protein structures produced by AlphaFold, Swiss-Model, I-TASSER, Phyre2, C-QUARK, and Robetta. The assessments include Ramachandran favored region percentages, ERRAT scores, VERIFY 3D scores, overall G-factors, planar group assessments, QMEAN scores, Global-Scores, and Z-scores. This comparative analysis offers a comprehensive understanding of the accuracy and reliability of each protein structure prediction tool [29, 37, 45, 67, 68]. Ensuring that the residues meet the expected structural criteria boosts confidence in the model’s predictions and its potential use in scientific research and development: For ROBEETA, 71.23% of the residues have an average 3D-1D score greater than or equal to 0.1. In AlphaFold, 69.86% of the residues meet this threshold. For SWISS-MODEL, 67.40% of the residues achieve an average 3D-1D score of 0.1 or higher. I-TASSER shows 68.49% of residues with scores meeting or exceeding 0.1. For Phyre2, 64.86% of the residues have a 3D-1D score of at least 0.1. Lastly, in C-QUARK, 65.75% of residues attain an average 3D-1D score of 0.1 or greater. These results indicate the proportion of residues in each model that meet the 3D-1D score threshold and indicate the overall quality and reliability of the protein structures generated by these different modeling tools [40, 41].

AlphaFold stands out as the top choice for the best protein structure model. Based on the average of key quality metrics, AlphaFold achieved the highest overall score of 34.17, demonstrating a reliable and well-rounded model across multiple aspects of structural quality. Its scores reflect excellence in several aspects: a Ramachandran favored percentage of 93.7% (Figure 2) indicates good stereochemistry [68], and a high ERRAT score of 98.9 highlights low error rates in the structure [69]. Additionally, AlphaFold shows strong VERIFY 3D compatibility (69.86) and an overall G-Factor of 0.22, further underscoring the reliability of its stereochemistry [37, 40]. Moreover, AlphaFold maintains planar group accuracy at 93.9%, supporting high-quality bond geometry [37]. Although the QMEAN score was not available in this dataset, prior analysis suggests that AlphaFold typically scores well in QMEAN (approximately 0.457).

**Figure 2.**
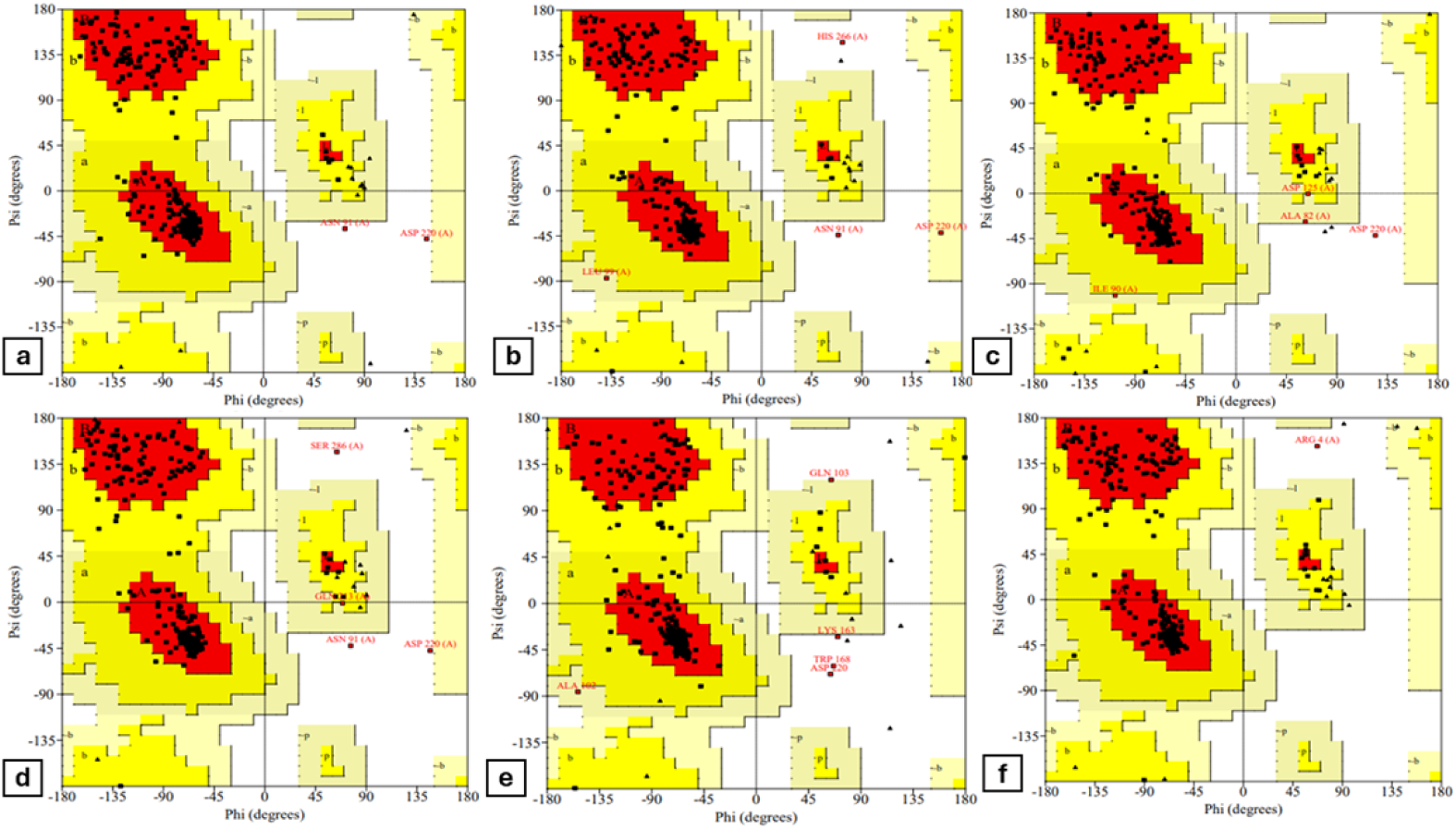
Ramachandran plots of (a) AlphaFold, (b) Swiss-Model, (c) I-Tasser, (d) Phyre2, (e) C-QUARK, and (f) Robetta that was obtained by SAVES server.

Other models, such as Phyre2 and Robetta, also perform exceptionally well. Phyre2 boasts the highest ERRAT score (99.6) among all models, indicating outstanding structural quality, coupled with a QMEAN score of 0.445 and a moderate Z-score of -7.98. However, its VERIFY 3D score (64.86) is slightly lower compared to AlphaFold. Robetta, on the other hand, excels with the highest VERIFY 3D score (71.23), indicating strong sequence-structure compatibility, and also achieves favorable G-factor (0.24) and QMEAN (0.453) scores.

### 3.3. Predicted Lddt and Aligned Error Heatmap of AlphaFold structure

The Predicted Aligned Error (PAE) chart shows the alignment errors between pairs of residues. Darker shades on the heat map indicate lower error values, signifying better alignment. The extensive dark green areas imply that the model anticipates low alignment errors for most residue pairs, bolstering the predicted structure’s accuracy [70]. The Predicted Local Distance Difference Test (PLddt) scores illustrate the confidence levels in the predicted positions of residues throughout the protein sequence, with higher scores denoting greater accuracy [70]. The result indicates that the majority of residues exhibit high pLDDT values, implying a strong confidence in the predicted structure of these regions (Figure 3) [71].

**Figure 3.**
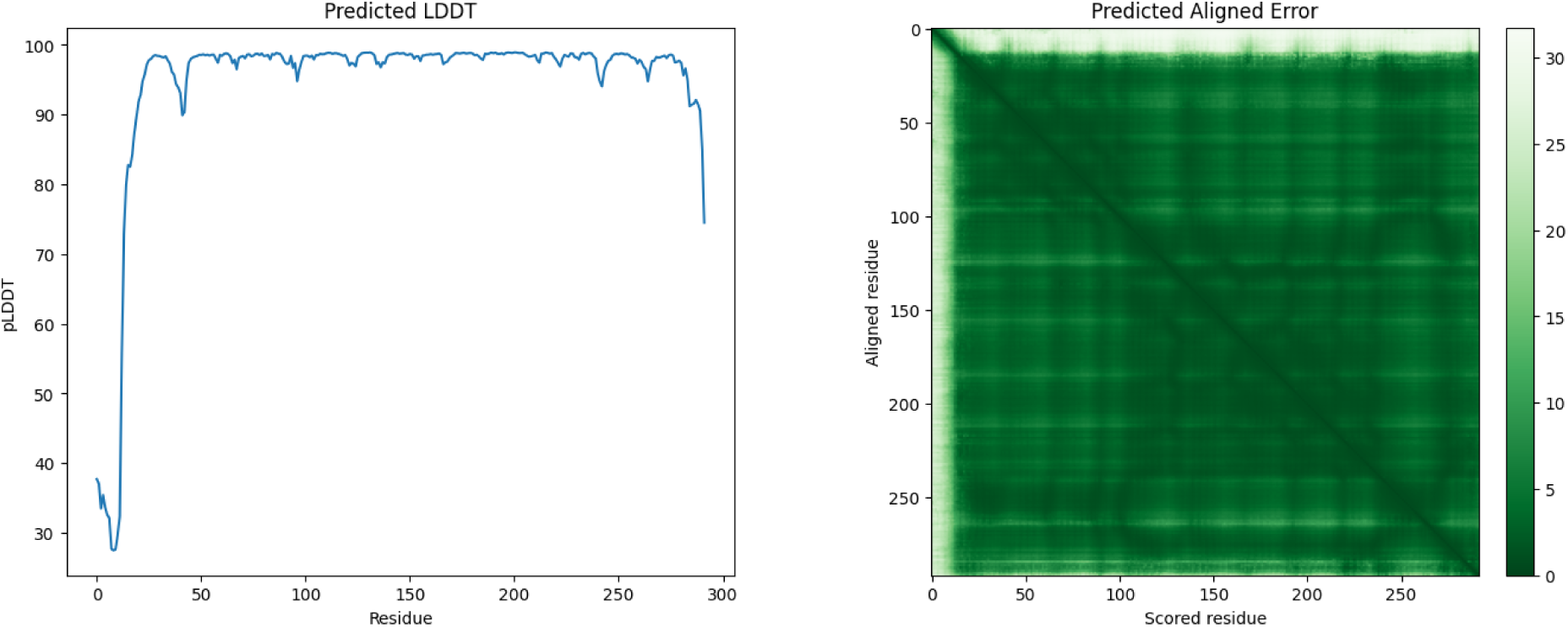
Predicted Local Distance Difference Test (pLDDT) and Predicted Aligned Error (PAE) for AlphaFold structure

Together, these visualizations offer a detailed evaluation of the protein structure’s reliability, showcasing regions of high confidence and minimal alignment error, which are crucial for precise protein function analysis and experimental validation. This high confidence is crucial for the reliability of the protein model, which is essential for comprehending its function and directing further experimental validation.

### 3.4. Molecular Docking

Molecular docking was carried out using AutoDock Vina integrated into the PYRx - Python Prescription 8.0 software. This powerful tool efficiently predicts the binding interactions between ligands and receptor sites. During the docking process, various possible conformations of the compounds were examined, and the most favorable binding conformation, which exhibited the highest binding affinity, was selected for visualization and further analysis. This method ensures that the optimal interactions are identified, providing a solid basis for subsequent experimental validation and functional studies. The compounds were ranked by their binding affinities in descending order (supplementary 1). CID 441035 has the highest binding affinity at -5.8 kcal/mol, followed by CID 439357 with -5.7 kcal/mol, and CID 840 with -5.6 kcal/mol. D-Ribose with CID 1097565, used as a control, also has a binding affinity of -5.3 kcal/mol.

Based on these results, CID 441035 related to D-Talopyranose, was selected for further analyses due to its strongest binding affinity. D-Talopyranose, also known as D-Talose, is a hexose sugar with the molecular formula C6H12O6 (Figure 4). This monosaccharide has a molecular weight of approximately 180.16 g/mol and is characterized by its cyclic pyranose form. It features multiple hydroxyl groups attached to its carbon atoms. D-Talose plays a role in various biological processes, including metabolism in certain organisms like bacterias and can be biosynthesized [72]. Based on these findings, CID 441035 was selected as the best candidate for further analyses due to its strongest binding affinity.

**Figure 4.**
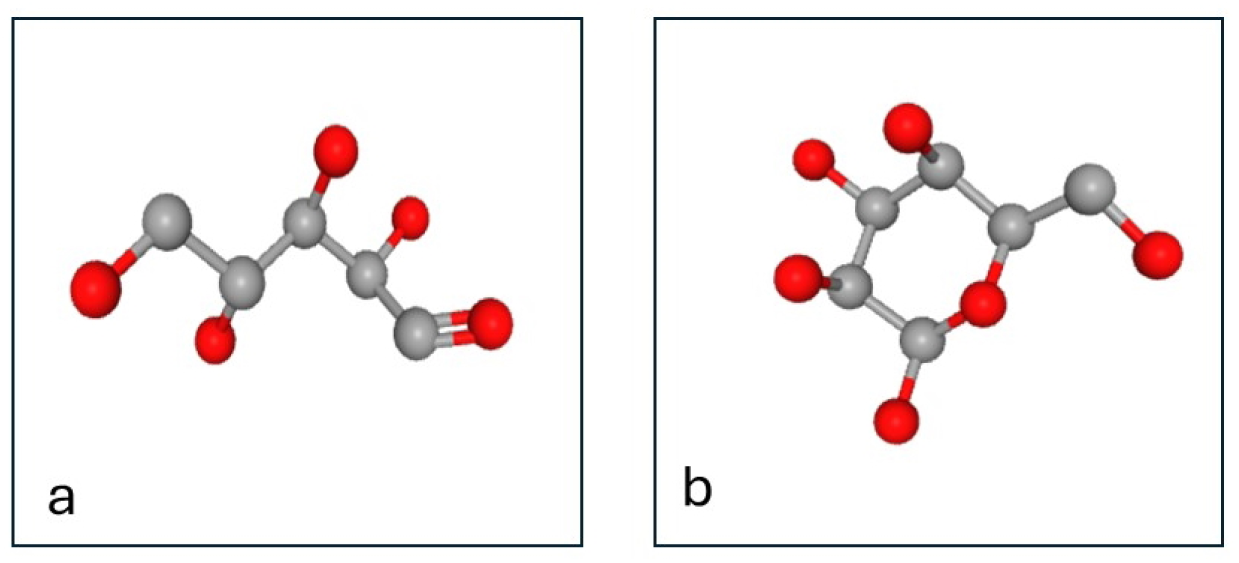
3D structures of a) D-Ribose and b) D-Talose without hydrogens

### 3.5. Visualization

Following the docking simulations, D-Talose interactions with the AlphaFold structure were visualized using Mol* server [73]. This powerful molecular visualization tool allowed for a detailed examination of the binding poses, offering critical insights into how the ligand interacts with the protein’s active site (Figure 5). These visualizations confirmed the predicted binding affinities and highlighted key molecular interactions, supporting the docking simulation results and aiding in the drug discovery process.

**Figure 5.**
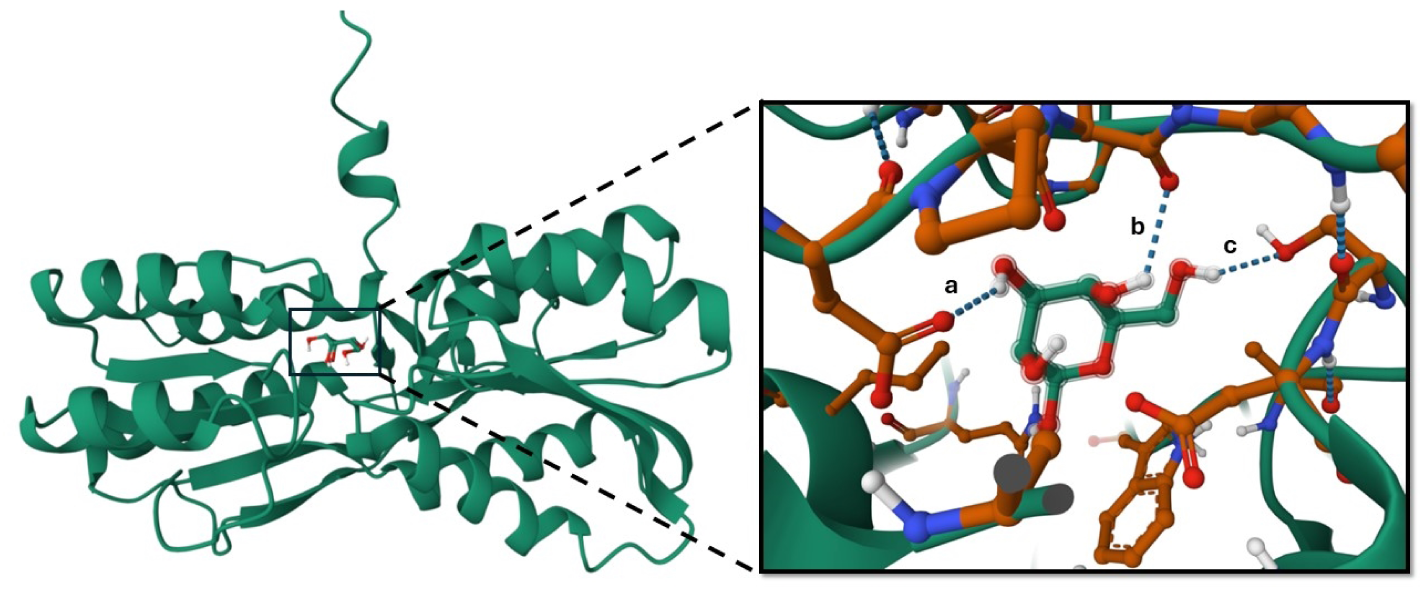
Visualization of the protein-ligand complex showing key hydrogen bonds between D-Talose and DBP in 5 Angstroms distance. visualization was performed in Mol* server which a, b, and c refers to hydrogen bonds with SER137, ALA68, ASP66

The visualization reveals that amino acids SER137, ALA68, ASP66 establish hydrogen bonds with the ligand at a distance of roughly 5 Ångström. This indicates specific interactions between these residues and the ligand, which are vital for the binding affinity and stability of the protein-ligand complex. These interactions offer insights into the binding mechanism and help elucidate the functional significance of the complex [73].

### 3.6. ADMET Analysis

To evaluate the ADMET characteristics of D-Talose, we utilized the related isomeric SMILES formula "C([C@@H]1[C@@H]([C@@H](<u>C@@H</u>O)O)O)O". The result provides a comprehensive analysis of various pharmacokinetic, pharmacodynamic, and toxicity parameters for the compound, highlighting its potential as a drug-like molecule (Figure 6). The physicochemical properties of the compound include a molecular weight of 180.06, which is within the optimal range for drug candidates. Key features such as the number of hydrogen bond donors (5) and acceptors (6), rotatable bonds (1), and a topological polar surface area (TPSA) of 110.38 are within favorable ranges, suggesting potential stability and compatibility with biological systems [55].

**Figure 6:**
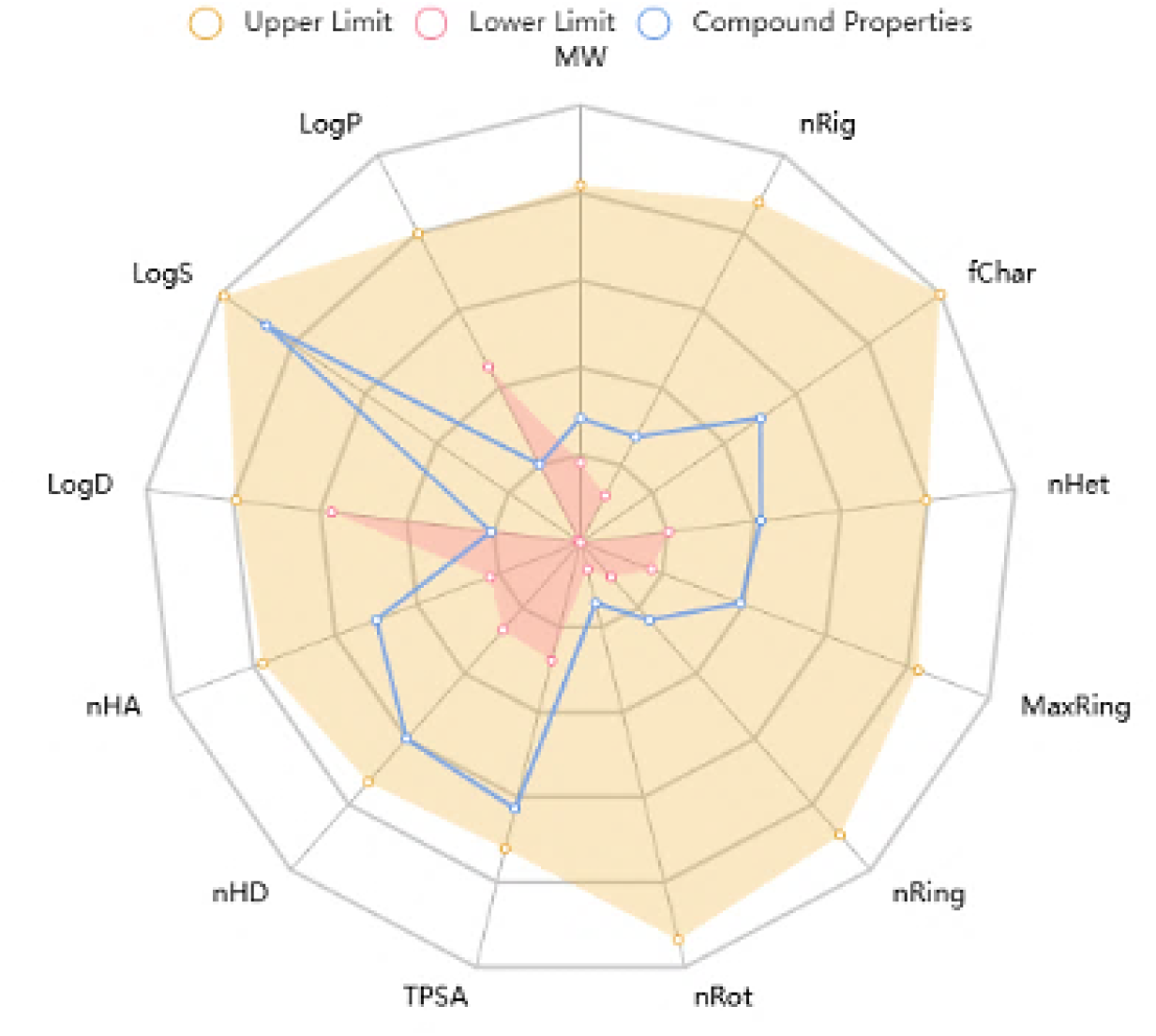
The radar chart from ADMETlab2 visualizes various properties of the compound. Each axis represents a different property, offering a comprehensive overview of the compound’s characteristics. These properties include LogP, which measures lipophilicity, and LogS, which indicates solubility in water. LogD reflects lipophilicity at a specific pH, while nHA (number of hydrogen bond acceptors) and nHD (number of hydrogen bond donors) are crucial for interactions with biological targets.

Its aqueous solubility (logS = −0.046) and partition coefficient (logP = −2.19) are also within optimal values (Figure 6), indicating balanced solubility and permeability, which are crucial for effective bioavailability. In medicinal chemistry assessments, the compound’s drug-likeness score (QED = 0.29) is on the lower end of desirability; however, it scores well in synthetic accessibility (SA score = 3.595), indicating it is relatively easy to synthesize. The compound meets significant drug-likeness criteria such as Lipinski, Pfizer, and GSK, which evaluate attributes like molecular weight, logP, and hydrogen bond characteristics. However, it does not meet the "Golden Triangle" criteria, which are often associated with favorable ADMET properties [74].

The absorption properties show mixed results, with low Caco-2 permeability (−5.384), indicating limited intestinal absorption. However, there is a moderate probability of human intestinal absorption (HIA = 0.848), suggesting a reasonable chance of bioavailability. Additionally, the compound has low probabilities of being a P-glycoprotein (Pgp) inhibitor (0.002) or substrate (0.054), potentially reducing the risk of efflux-related absorption issues. The distribution profile appears favorable, with plasma protein binding (PPB) being relatively low at 12.88%. Additionally, the compound exhibits a high unbound fraction of 78.7%, indicating that it will be readily available in the plasma for active distribution. The compound also shows a low probability of crossing the blood-brain barrier (BBB penetration = 0.319), suggesting it may not be effective for central nervous system (CNS) targets but could be suitable for peripheral applications.

The metabolism data indicates that D-Talose does not appear to intervene with key cytochrome P450 (CYP) enzymes like CYP1A2, CYP2C19, CYP2C9, CYP2D6, and CYP3A4. This is a superb sign as it approaches the compound is not likely to cause serious drug-drug interactions, that is truely beneficial for its use as a remedy. In terms of excretion, it has a mild clearance fee of 1.492 mL/min/kg and a quick half of-lifestyles of 0.816 hours. Basically, this indicates the compound is quick cleared from the body, reducing the danger of it constructing up to harmful tiers. However, as it leaves the body so fast, it would be better to take it more regularly to keep the medicine powerful.

Toxicity assessments indicate generally low toxicity risks for the compound. It has a low probability of being a hERG inhibitor (0.068), reducing the risk of cardiotoxicity. Additionally, the compound shows low scores for human hepatotoxicity (0.044) and drug-induced liver injury, as well as other toxicity measures such as skin sensitization and acute toxicity, suggesting it is potentially safe for clinical use. However, environmental toxicity assessments raise some concerns. The compound shows alerts for aquatic toxicity and non-biodegradability, indicating potential adverse environmental impacts if introduced in significant amounts into ecosystems. Overall, the compound exhibits several promising drug-like qualities, particularly in terms of ease of synthesis, distribution, and low toxicity.

Additionally, the chart displays TPSA, impacting absorption and permeability, and nRot (number of rotatable bonds), indicating molecular flexibility. Other properties, such as nRing (number of rings), MaxRing (maximum ring size), and nHet (number of heteroatoms), provide further insights into the compound’s structural features. Formal charge (fChar) and the number of rigid bonds (nRig) are also shown, contributing to an understanding of the compound’s stability and interaction potential. The radar chart features three types of data points: the upper limit (yellow circles), the lower limit (pink circles), and the compound properties (blue line). This visual representation helps compare the compound’s properties against predefined limits, aiding in assessing its suitability for further development or study.

The radar chart visually represents the compound’s properties against drug-likeness thresholds, offering an overview of its alignment with ideal pharmacokinetic and structural criteria. Starting with molecular weight (MW), the compound comfortably fits within the upper and lower limits, indicating compliance with typical drug standards. The LogP and LogS values, reflecting the compound’s lipophilicity and solubility, are also within the desired range. However, the LogS value is near the lower boundary, suggesting that although the compound is somewhat soluble, it is close to the limit of acceptable solubility. Similarly, the LogD value, which measures lipophilicity at physiological pH, lies within drug-like boundaries, indicating that the compound effectively balances solubility and permeability.

Both the number of hydrogen bond acceptors (nHA) and donors (nHD) for the compound fall within favorable ranges, which are essential for interactions with biological targets and contribute to its potential bioactivity. The topological polar surface area (TPSA), impacting absorption and permeability, also falls well within the desired range, suggesting the compound’s suitability for oral bioavailability. The number of rotatable bonds (nRot), which affects molecular flexibility, is within acceptable limits, indicating good bioavailability and ease of absorption. Additionally, other structural features, such as formal charge (fChar), number of heteroatoms (nHet), maximum ring size (MaxRing), and number of rings (nRing), meet the drug-likeness criteria, highlighting that the compound’s molecular framework is well-suited for biological activity. Lastly, the number of rigid bonds (nRig) is within the acceptable range, contributing to a stable molecular structure that supports favorable pharmacokinetics.

Overall, the blue line depicting the compound’s properties mostly remains within the orange boundary (upper limit) and outside the red area (lower limit). This suggests that the compound has a well-balanced set of drug-like properties. However, certain aspects, such as solubility, might need further optimization based on its intended application.

### 3.7. Molecular Dynamics

Molecular dynamics simulations for the target protein and its top protein-ligand complexes were performed using GROMACS 2021.4. The initial protein structure was prepared using the CHARMM27 force field and the TIP3P water model. The complex was placed in a triclinic box with a 1.0 nm buffer and solvated with the SPC216 water model. To ensure neutrality, Na^+^ and Cl^−^ ions were added to reach a 0.1 M concentration. Energy minimization was conducted to remove unfavorable contacts, followed by equilibration in two phases: NVT (constant Number, Volume, Temperature) and NPT (constant Number, Pressure, Temperature).

After the simulation, the trajectory data was re-centered and re-wrapped to ensure precise analysis. To evaluate the stability of the protein-ligand complexes, several stability and interaction analyses were performed, including RMSD, RMSF, hydrogen bond count, radius of gyration, and energy calculations. Xmgrace was used for visualization, enabling a detailed assessment of binding affinities and molecular interactions.

The Root Mean Square Deviation (RMSD) of the protein’s backbone (Figure 7) increased throughout the 5-nanosecond simulation, indicating that the protein’s structure deviates from its initial conformation over time. This suggests alterations in the protein’s stability and conformational dynamics during the molecular dynamics simulation, which is crucial for understanding the protein’s behavior in a simulated environment and evaluating the stability and interactions of protein-ligand complexes. the RMSD increases and fluctuates but does not show a large, continuous increase, suggesting that the protein backbone has reached a relatively stable state, though with some conformational flexibility.

**Figure 7:**
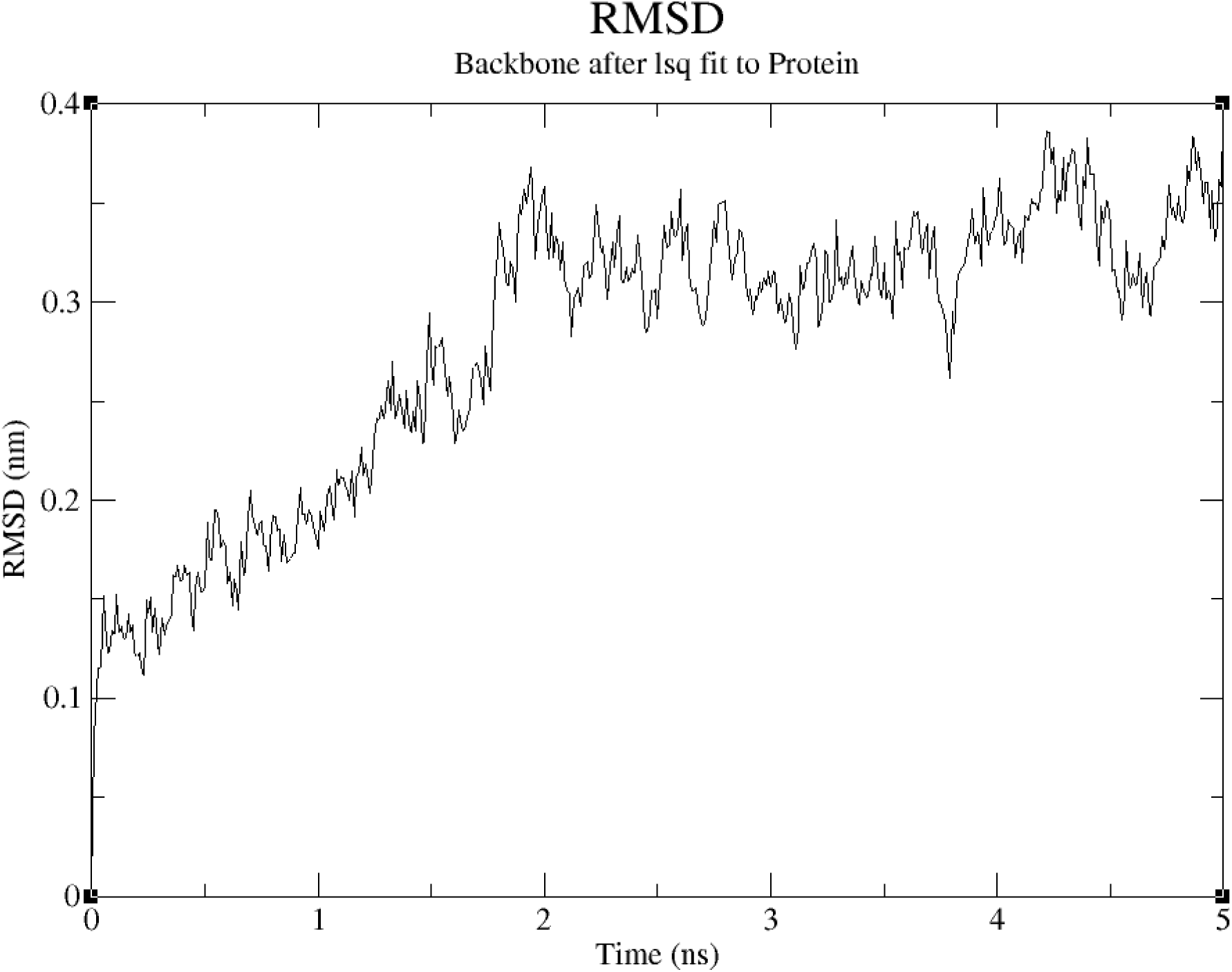
The RMSD (Root Mean Square Deviation) of protein backbone

The RMSF assesses the flexibility of each atom in the protein structure (Figure 8). Initially, RMSF values are around 1 nm, stabilizing at approximately 0.2 nm for most atom indices as the simulation progresses, with a slight increase towards the end. This suggests that while most protein atoms remain stable throughout the simulation, some regions exhibit higher flexibility that can be in active site.

**Figure 8:**
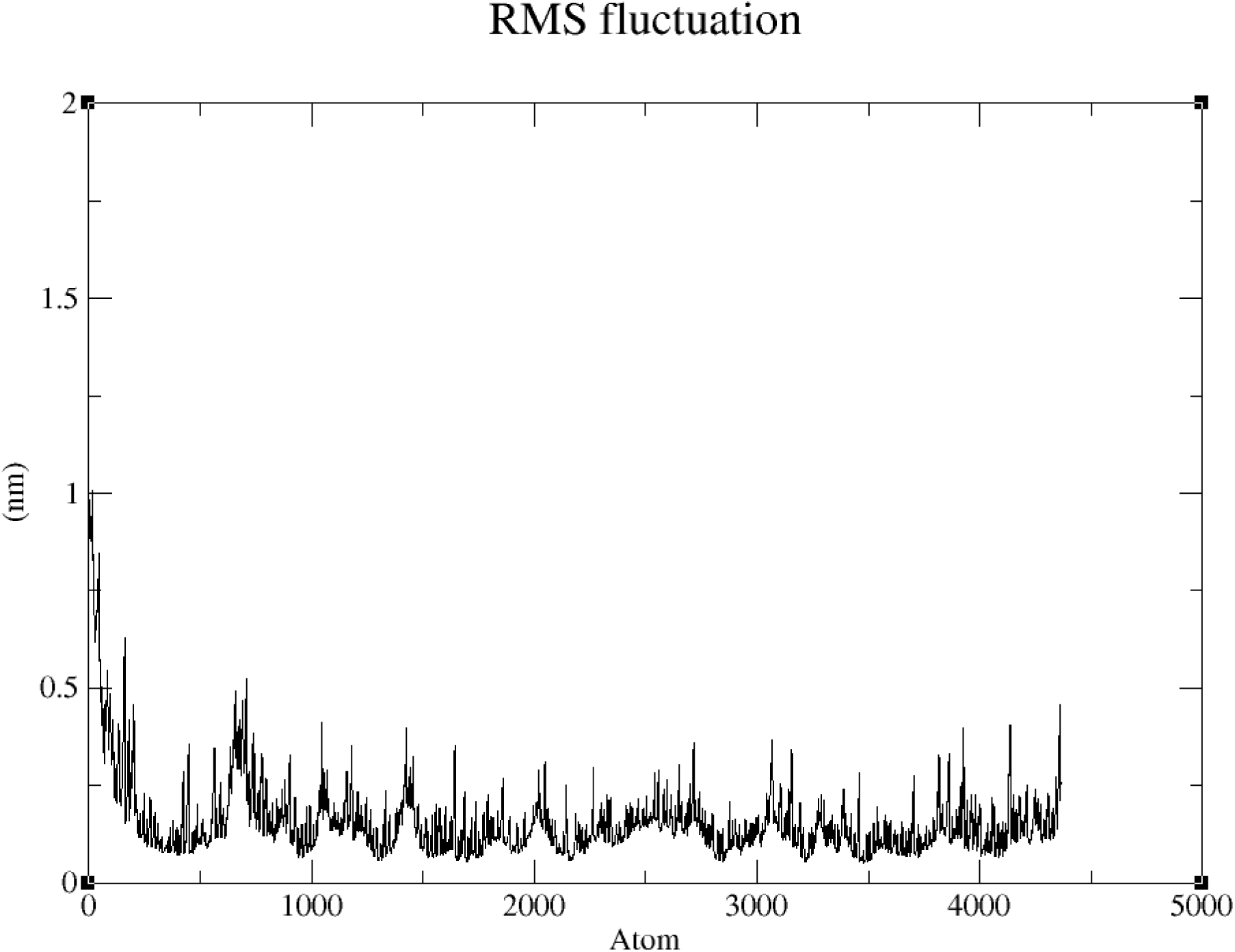
The RMSF (root mean square fluctuation) of protein structure.

The overall Radius of Gyration (Rg) values (Figure 9) remain relatively stable throughout the simulation, indicating that the protein retains its compactness over time. This suggests that the protein’s structure does not significantly expand or contract, reflecting its consistency during the molecular dynamics simulation. However, when examining the Rg along specific axes (x, y, and z), some fluctuations are observed, highlighting minor structural changes and reorientations within the protein that provide insights into how different parts of the protein move and adjust during the simulation.

**Figure 9:**
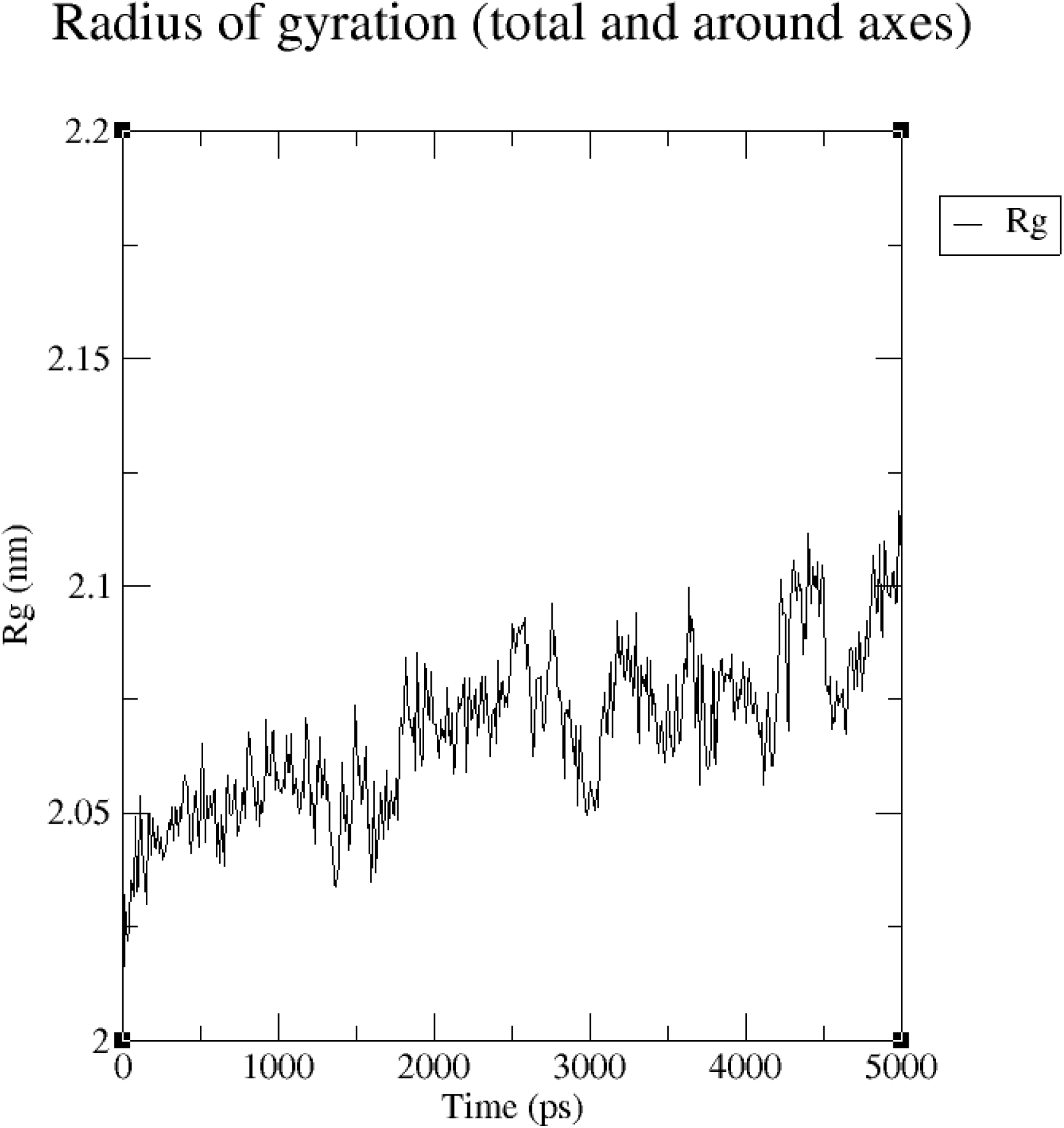
The overall Radius of Gyration (Rg) indicating that the protein compactness over time.

The decreasing trend in the number of hydrogen bonds (Figure 10) indicates that the stability and interactions within the protein-ligand complex lessen as the simulation progresses. This suggests a potential loss of structural integrity or changes in bonding patterns, which are crucial for understanding the dynamics and stability of the complex. Further analysis of specific regions and residues involved in these hydrogen bonds could provide deeper insights into the structural changes occurring within the complex.

**Figure 10:**
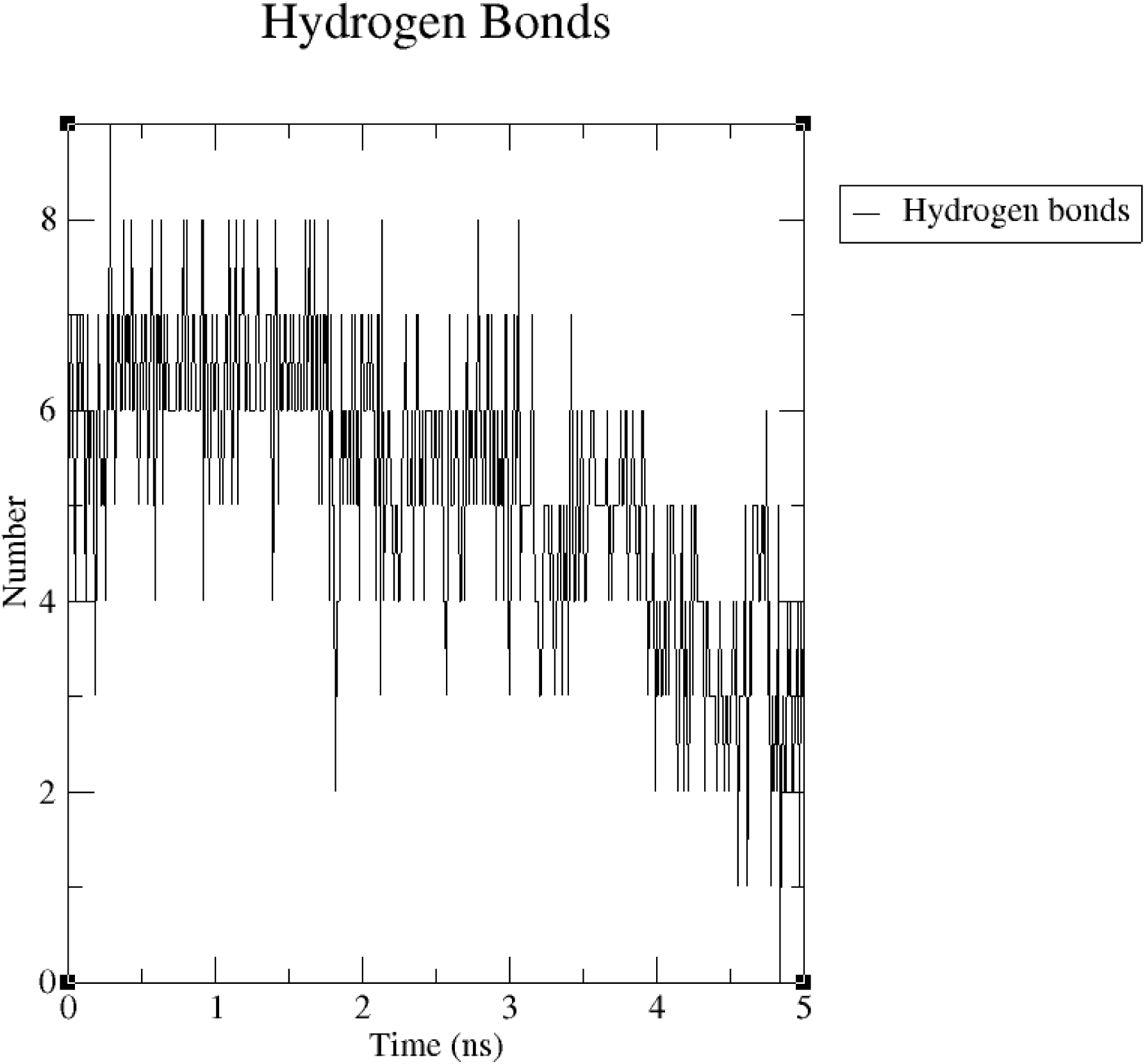
number of hydrogen bonds within the protein-ligand complex

The potential energy of the system, ranging from 0 to 5,000 picoseconds (Figure 11), shows fluctuations between approximately −618,000 and −612,000 kJ/mol, reflecting the dynamic nature of molecular interactions within the system throughout the simulation period. The consistent range of potential energy values suggests that the system maintains a relatively stable state over the course of the simulation. This stability is essential for understanding the energetic behavior and structural integrity of the molecular system during the GROMACS simulation run, providing crucial insights into the system’s molecular dynamics and overall stability.

**Figure 11:**
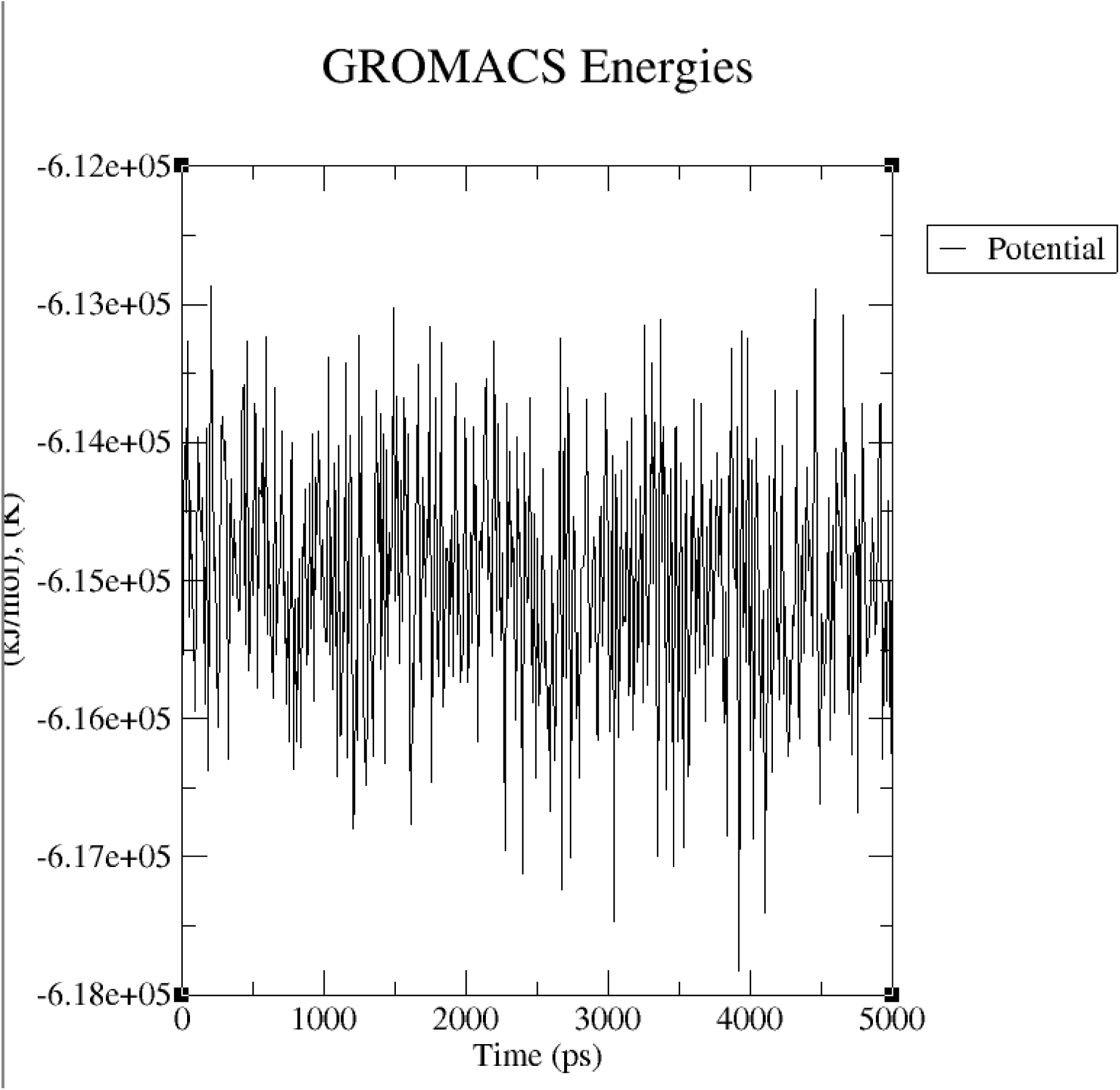
The potential energy of the system

## 4. Conclusion

This study underscores the potential of targeting the D-Ribose Binding Protein (DBP) in *Brucella melitensis* as an innovative therapeutic strategy for brucellosis. By disrupting this protein, essential for nutrient uptake and bacterial metabolism, we can inhibit Brucella’s growth and pathogenicity, presenting an alternative to conventional antibiotic treatments. Through structural modeling, molecular docking, and molecular dynamics simulations, D-Talopyranose emerged as a promising ligand with strong binding affinity and favorable ADMET properties, suggesting its suitability as a drug candidate. Given the rising challenge of antibiotic resistance, this approach highlights a novel pathway for brucellosis treatment, emphasizing the importance of developing specific inhibitors for bacterial proteins integral to survival and infection. Further experimental validation and optimization of these inhibitors could pave the way for effective, targeted therapies against Brucella, contributing to improve disease management and public health outcomes.

## 5. Suggestions

To further develop D-Talopyranose as a therapeutic candidate against *Brucella melitensis*, several important steps are proposed. First, experimental validation through in vitro and in vivo studies is crucial to confirm the compound’s binding efficacy with the DBP. These studies will provide essential insights into D-Talopyranose’s inhibitory potential and effectiveness as a therapeutic agent. Additionally, optimizing the ligand structure through modifications aimed at enhancing D-Talopyranose’s binding affinity, stability, and selectivity could improve its therapeutic properties. Structure-activity relationship (SAR) studies would be valuable in refining the molecule’s characteristics for increased drug efficacy. To advance the drug discovery process, further validation of binding affinity, ADMET profiling, and molecular dynamics simulations should be performed on the compound with the highest binding affinities, namely CID 439357 and 840. Additionally, optimizing the structures of these compounds could enhance their binding interactions and specificity, potentially leading to new and effective therapeutic agents for brucellosis.

